# A systematic structural comparison of all solved small proteins (4-6 kDa) reveals the weight of disulfide bonds in proteins’ foldability

**DOI:** 10.1101/2021.03.30.437752

**Authors:** Mariana H. Moreira, Fabio C. L. Almeida, Tatiana Domitrovic, Fernando L. Palhano

**Affiliations:** Programa de Biologia Estrutural, Instituto de Bioquímica Médica Leopoldo de Meis, Universidade Federal do Rio de Janeiro, Rio de Janeiro, RJ, 21941-902, Brazil; Departamento de Virologia, Instituto de Microbiologia Paulo de Góes, Universidade Federal do Rio de Janeiro, Rio de Janeiro, 21941-902, Brazil

## Abstract

Defensins are small proteins, usually ranging from 4 to 6 kDa, amphipathic, disulfide-rich, and with a small or even absent hydrophobic core. Since a hydrophobic core is generally found in globular proteins that fold in an aqueous solvent, the peculiar fold of defensins can challenge tertiary protein structure predictors. We performed a PDB-wide survey of small proteins (4-6 kDa) to understand the similarities of defensins with other small disulfide-rich proteins. We found no differences when we compared defensins with non-defensins regarding the proportion and exposition to the solvent of apolar, polar, and charged residues. Then we divided all small proteins (4-6 kDa) deposited in PDB into two groups, one group with at least one disulfide bond (bonded, defensins included) and another group without any disulfide bond (unbonded). The group of bonded proteins presented apolar residues more exposed to the solvent than the unbonded group. The *ab initio* algorithm for tertiary protein structure prediction Robetta was more accurate to predict unbonded than bonded proteins. Our work highlights one more layer of complexity for the tertiary protein prediction structure: small disulfide-rich proteins’ ability to fold even with a poor hydrophobic core.

## Introduction

Defensins are a group of small proteins (< 10 kDa), their primary sequence is very diverse but rich in cysteine, and their function is related to host defense(1, 2). Defensins can be found in animals, plants, and fungi(2, 3). In terms of structure, defensins proteins form disulfide bonds, which seem to stabilize their tertiary structure(1, 2, 4, 5). Defensins present a large variety of primary sequences. Still, the tertiary structure has a compact core, and they typically show a triple-stranded antiparallel β-sheet, packed against an α-helix restrained by disulfide bonds(2). Defensins can afford an unusual structure lacking a hydrophobic core, with a high proportion of hydrophobic residues exposed to the solvent(5, 6)}. Other globular proteins form a hydrophobic core, which buries apolar residues, minimizing their solvent accessibility surface(7). In 2018, Almeida and collaborators solved Sugarcane defensin 5 and *Pisum sativum* defensin 1. Both proteins also lack a hydrophobic core and presented an unusual side-chain exposition of multiple hydrophobic amino acids(8). They concluded that defensins are stabilized by tertiary contacts formed by surface-exposed hydrophilic and hydrophobic residues(8). Later, in 2020, Almeida’s group solved the structure of *Pisum sativum* defensin 2, and they noticed the same unusual fold(9). It was hypothesized that the long polar/charged side chains at the surface-clusters protect the hydrophobic amino acids from complete exposure to the solvent(9). The features that allow the poor hydrophobic core of defensins to exist are still to be determined, but likely the presence of disulfide bonds plays an essential role in their fold.

Even though it is well known that defensins possess an unusual fold regarding the poor hydrophobic core, as far as we know, no systematic comparison of defensins with other small proteins containing or not disulfide bonds was performed yet. We analyzed all small protein structures (4-6 kDa) with at least one disulfide bond deposited on Protein Data Bank (PDB) to address this issue. The parameters we chose to compare defensins vs. non-defensins proteins were the proportion and the degree of solvent exposition of apolar, polar, and charged residues. No statistical significance was found for these analyses suggesting that other small proteins that form disulfide bonds besides defensins also present an unusual fold lacking a canonical hydrophobic core. Next, we compared all PDB deposited small proteins (4-6 kDa) containing (bonded) or not (unbonded) disulfide bonds following the same rationale. This time, we observed that the bonded group has a lower proportion of apolar residues than the unbonded group. Those apolar residues were more exposed to the solvent, compromising their hydrophobic cores consequently. We also compared the frequency and exposition of the long polar/charged side chains between the two groups, but no differences were found. With the recent progress of deep learning applied to protein structure determination, this field has significantly improved. Nevertheless, prediction servers still have a long way to go (10–13). We challenge the *ab initio* algorithm Robetta, which predicts tertiary protein structure, comparing the accuracy of prediction between bonded vs. unbonded peptides. Interestingly, the Robetta accuracy was higher for the unbonded peptides vs. bonded since the former possess the “canonical” hydrophobic core. Our study points out that even for small proteins, the prediction algorithms still have difficulty determining the protein structure, especially if the protein forms disulfide bonds.

## Materials and Methods

### Data collection and analysis workflow

To understand how defensins are comparable to other small proteins, we chose the molecular weight ranging from 4 to 6 kDa because it allowed us to obtain a significant proportion of defensins in the bonded group (with disulfide bond). At the same time, submit data under a meticulously manual review. Figure 1 shows a visual representation of the data collection and refinement until results analysis. We performed an advanced search of PDB files in the RCSB PDB database and used the Uniprot database to collect FASTA sequences of the selected proteins. In advanced search options of the RCSB PDB database, we could not refine as much as we aimed to obtain only proteins in an aqueous solvent, without ligands or other elements that could interfere in their folding, so we had to cure the data manually. After manual review, the groups were significantly reduced because we aimed to obtain unbiased comparable groups in which the proteins do not have any variable that would interfere in their folding in an aqueous solvent. Bonded and unbonded groups are proteins with and without at least one disulfide bond.

**Figure 1.**
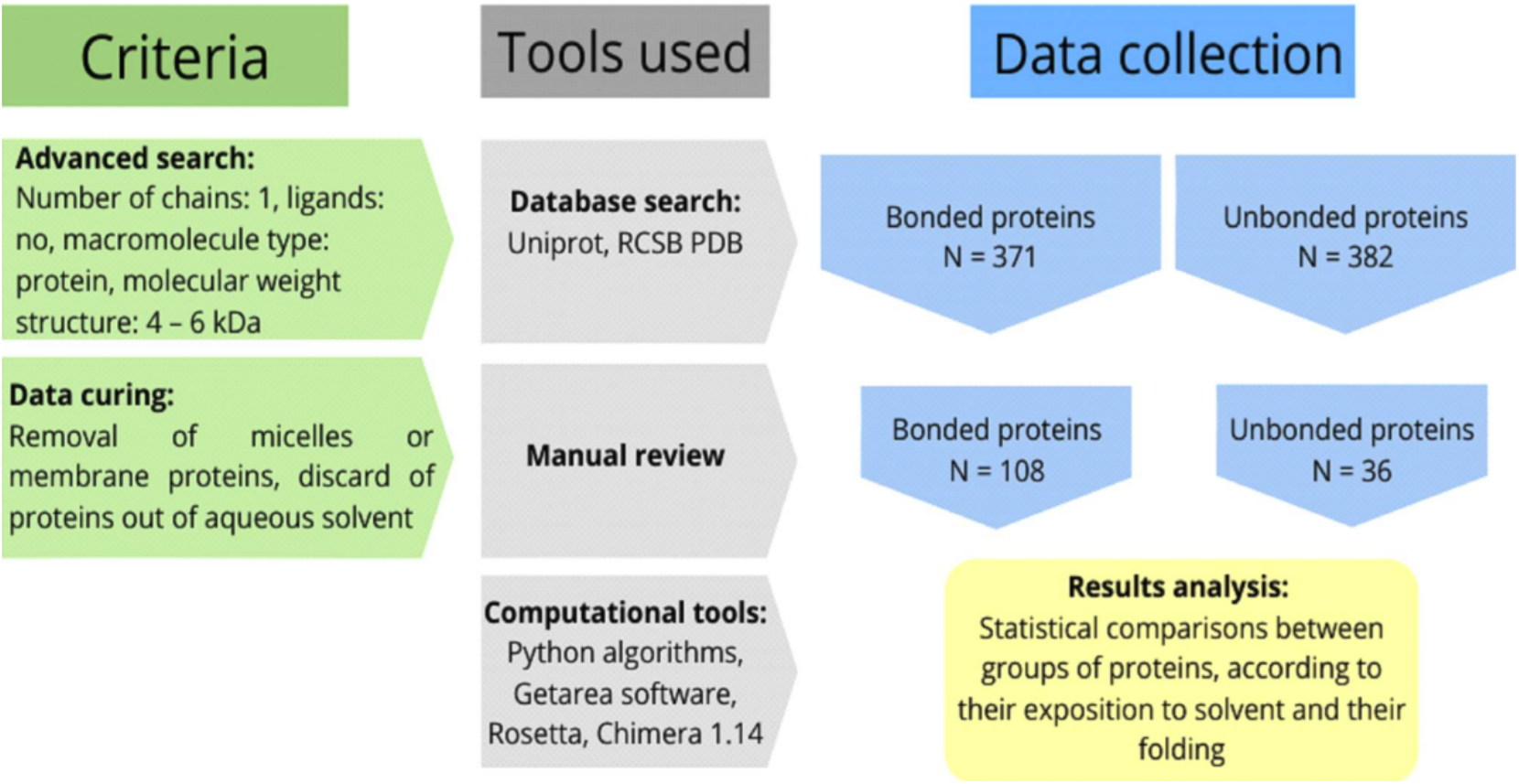
Schematic representation of the pipeline used herein to obtain curated groups of bonded and unbonded small proteins. After curing, 108 proteins for the bonded group and 36 proteins for the unbonded groups were selected.

### Protein curing

The database RCSB PDB available at rcsb.org was used as a source of 3D protein structures. Selected data were downloaded as a PDB file and submitted to the following analyses. In March 2020, advanced search options (number of chains: 1 to 1 AND ligands: no AND molecular weight: 4000 to 6000 AND disulfide bond: 0 to 9999 AND macromolecule type: protein only AND NOT structure description: micelle OR membrane) were used to obtain our data. In this first search, 753 structures were found. A second, advanced search changing the option “disulfide bond”: 1 to 9999 found 371 proteins. In this case, these proteins had at least one disulfide bond in their structure, so the first search had 382 proteins that did not have a disulfide bond at all.

Two groups were formed, one containing proteins with at least one disulfide bond, named bonded group, and the other containing proteins without disulfide bond, which was named unbonded group. Before analyzing this data, both groups were submitted to manual curing. Only proteins in an aqueous solvent, with a well-defined secondary structure and following all the criteria described in advanced search options were kept. Finally, the bonded group contained 108 proteins, and the unbonded group had 36 proteins. All data information, including analysis results, is in a supplementary table1.

### Solvent accessible surface area (SASA) calculation

To calculate SASA for each PDB file, we submitted them to the software GETAREA (14), online available at curie.utmb.edu/getarea.html, and the software Chimera 1.14(15, 16). The SASA result obtained by GETAREA is given in percentual of residue exposition, so the calculation by Chimera had to be transformed into a percentage result to be comparable with the other method.

After opening a PDB file in Chimera, the option “show” in “Surface” was selected from the “Action” menu. Then, using “Render by Attribute” from “Structure Analysis” in the “Tools” menu, the attributes “AreaSAS” and “AreaSES” of residues were exported in a TXT file. These files were submitted in a Python software that we developed called “SASA_chimera_calculation.py.” This algorithm calculates for each residue its percentage exposition by the following eq. 1:

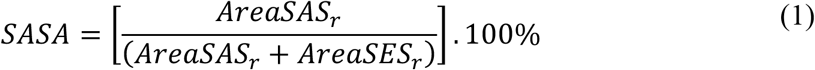

AreaSAS_r_ and AreaSES_r_ refer to each amino acid residue’s values in Chimera’s respective TXT files.

The percentage results obtained from both software are submitted to further Python programs to calculate the medium values of amino acid proportion and exposition.

### Amino acids proportion and exposition calculation

A Python software called percentage_proportion.py was developed to receive the results obtained by GETAREA and Chimera, and it classifies the amino acids according to Table 2. The percentage proportion of apolar, polar, and charged amino acids present in the structure is calculated for each protein.

**Table 1.**
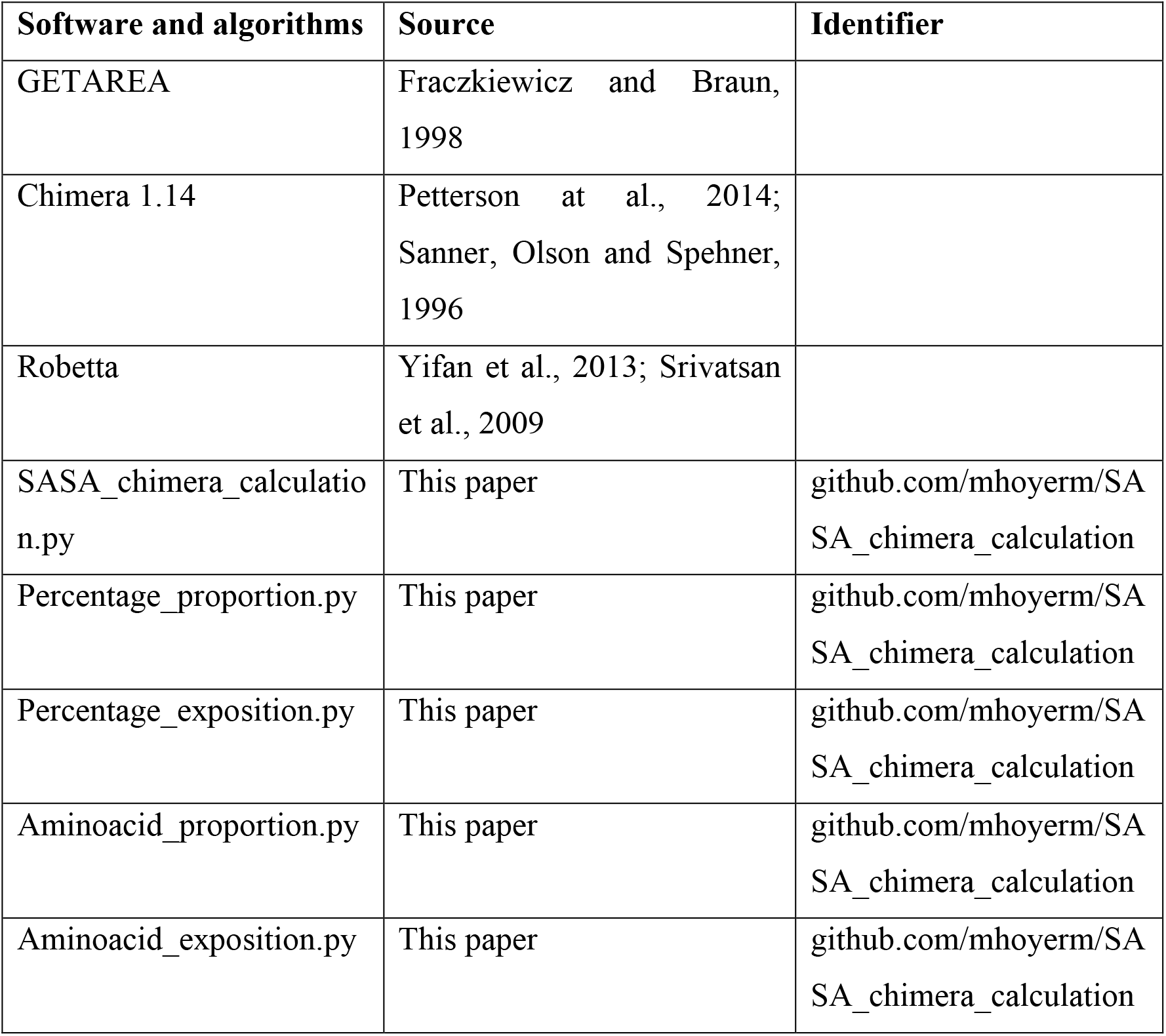
Software and algorithms used for data acquisition.

**Table 2.** Amino acid classification in apolar, polar, or charged.

| **Classification** | **Amino acids** |
| --- | --- |
| Apolar | GLY, ALA, VAL, PRO, LEU, ILE, MET, TRP, PHE |
| Polar | CYS, SER, THR, TYR, ASN, GLN |
| Charged | LYS, ARG, HIS, ASP, GLU |

Also, because the input files also contained SASA values of each amino acid, a similar Python software named percentage_exposition.py calculates the average exposition of apolar, polar, or charged residues. For each group of classified amino acids, the arithmetic mean of their exposition values is calculated. Thus, these Python software’s output files include percentage values of both amino acid proportion or average exposition in apolar, polar, and charged groups of amino acid residues present in the protein.

The software aminoacid_proportion.py aminoacid_exposition.py analyzed the tendency to form surface clusters with polar or charged amino acids. These algorithms receive the same inputs as the previous, but in this case, the amino acid classification follows Table 3(9). Like the earlier algorithms, this Python software’s output files include percentage values of amino acid proportion or average exposition, respectively, for amino acid residues classified as a short-chain or long-chain present in the protein.

**Table 3.** Amino acid classification of polar or charged amino acids with a short or long side chain.

| **Classification** | **Amino acids** |
| --- | --- |
| Short side chain | CYS, SER, ASN, ASP |
| Long side chain | THR, TYR, GLN, LYS, ARG, HIS, GLU |
| Non classified | GLY, ALA, VAL, PRO, LEU, ILE, MET, TRP, PHE |

### Ab initio 3D determination

The online server Robetta was used to predict our data’s ab initio protein structures(17, 18). Primary sequences were obtained from RCSB PDB or the database SwissProt from uniport.org. The primary protein sequence was submitted as a job for structure prediction in Robetta, with the option “AB only” selected. After determining the tertiary structure, the server provides five models, downloaded as a PDB file.

### RMSD calculation

The root-mean-square deviation (RMSD) between the protein structure deposited on RCSB PDB, and the protein structure obtained from Robetta, was calculated by Chimera 1.14. Both comparable structures were open in Chimera, then in the menu “Tools,” the option “MatchMaker” from “Structure Comparison” was used to select a reference structure, which would be the one downloaded from RCSB PDB, and a structure to match, which would be the one obtained from Robetta.

In cases where the reference contained multiple structures, only the first was considered. It was combined with all five possible matches (because Robetta provides five structure models) to determine the lowest RMSD value. The chain pairing selected option was “Best-aligning pair of chains between reference and match structure.” The alignment algorithm was Needleman-Wunsch, BLOSUM-62, 1 gap extension penalty. Options “include secondary structure score (30%)” and “compute secondary structure assignments” were selected. The option “iterate by pruning long atom pairs until no pair exceeds 2 angstroms” was chosen in matching.

### Statistical analyses, correlation, and raw data

The raw data used to create all Figures in this paper is available in the Supplementary Table. All statistical analyses were performed by GraphPad Prism 7. For Figure 1, the Kruskal-Wallis test was used. For Figure 3, Figure 5, and supplementary Figure 1, the Mann-Whitney test was used. For Figure 4, linear regression with a 95% confidence level was performed.

## Results and discussion

### Compared to other small cysteine-rich proteins, defensins do not appear to have an unusual fold regarding the poor hydrophobic core

To understand how the tertiary structure of defensines differs from other small disulfide proteins, we downloaded all PDB ranging from 4 to 6 kDa from RCSB PDB with at least one disulfide bond (bonded, 371 protein structures, Figure 1). After manual review, the proteins solved in the presence of ligands, micelles, detergents, lipids, organic solvents were eliminated, resulting in 108 proteins representing 29% of the initial sample (Figure 1). The main classes of proteins in this group were defensins and toxins, representing each class 30% of the total (Figure S1). Using the software GETAREA we measured the proportion and degree of exposition to the solvent of all amino acids’ side chains in those small proteins. First, we determined if, within the bonded group, defensines had to be analyzed separately. As shown in Figure 2, we divided the bonded group into three groups. One was composed of all small proteins (4-6 kDa) of PDB containing at least one disulfide bond (108 proteins, bounded). The first group possesses defensins and non-defensins proteins. Another group with just non-defensins proteins (75 proteins, bonded minus defensins), and the third group was composed only of defensins (33 proteins, defensines). These groups were analyzed for parameters such as amino acid classification (apolar, polar, and charged, Table 2) in terms of proportion within the primary sequence (Figures 2A-2C) and their exposition to the solvent (GETAREA analyzes, Figures 2D-F). However, Figure 2 shows that non-significant differences were found among the three groups of proteins. We then decided to proceed with the entire bonded group, N = 108, for all subsequent analyses.

**Figure 2.**
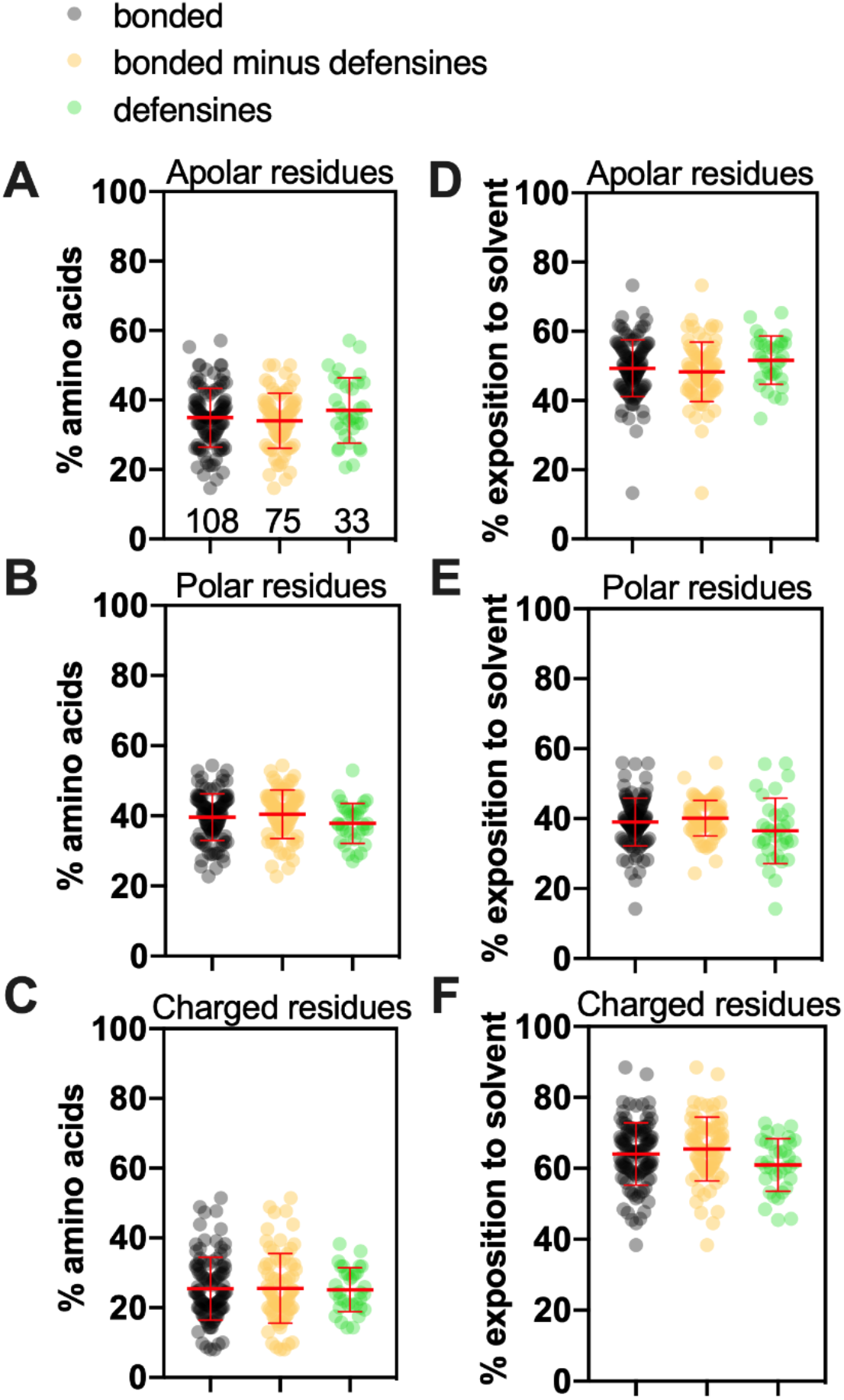
Defensines present a similar proportion and solvent exposition of apolar, polar, and charged residues compared to other small disulfide-bonded proteins. In all graphs, the whole population of bonded proteins is represented in black, N=108, the bonded group’s proteins without defensins are represented in orange, N=75, and the group containing only defensins is represented in green, N=33. The proportion of apolar (A), polar (B), or charged residues (C) was calculated for each protein. The average degree of solvent exposition was calculated using GETAREA (please see details in methodology). Exposition of apolar (D), polar (E), or charged (F) residues. Kruskal-Wallis test was performed for all comparisons, but no statistical differences were identified. Since no differences were observed, the following analyses involving the bonded group N = 108.

**Figure 3.**
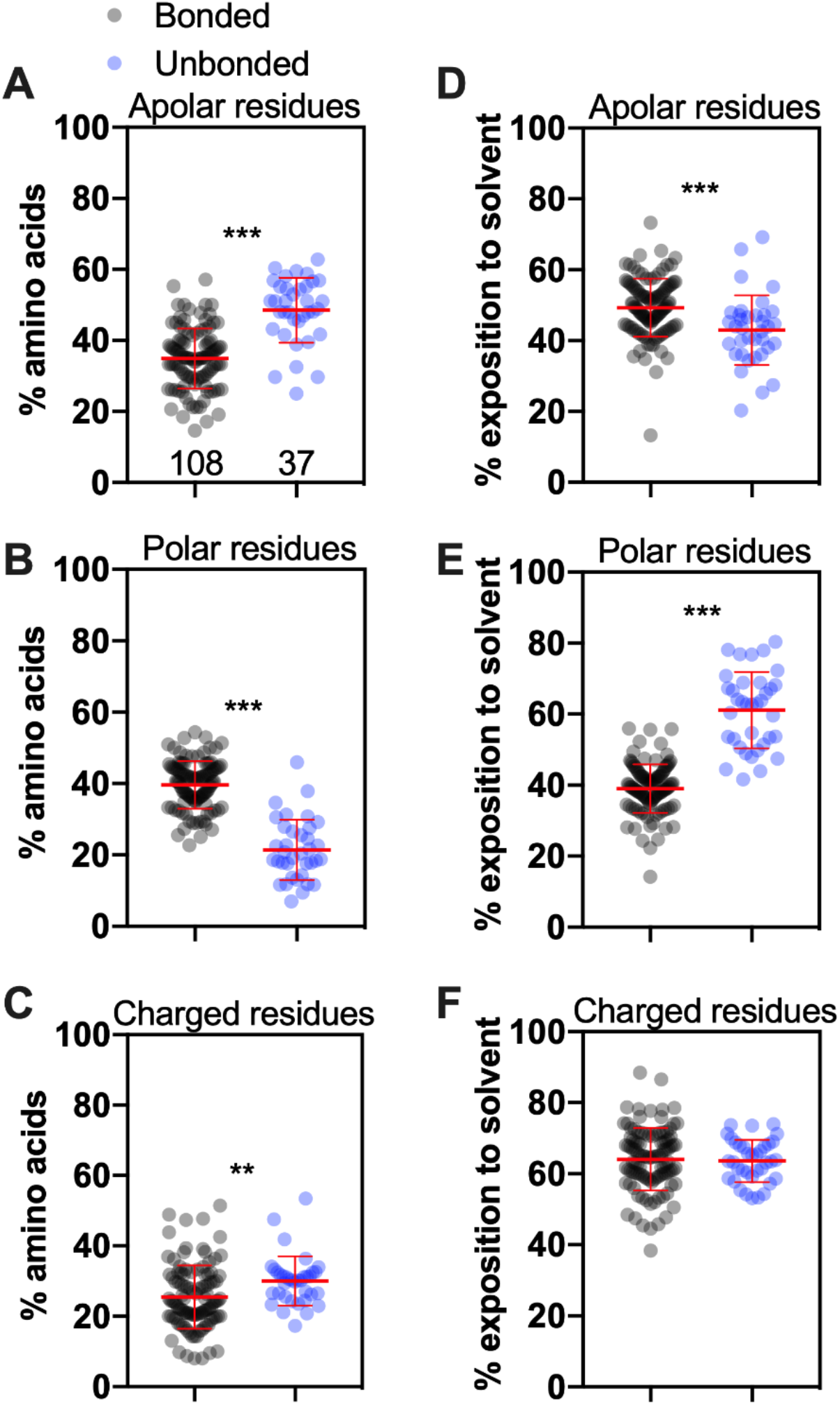
Small proteins with at least one disulfide bond (bonded) present distinct features regarding the proportion and exposition of residues compared to proteins that do not make disulfide bonds (unbonded). In all graphs, the whole population of bonded proteins is represented in black, N=108, while the group of unbonded proteins is represented in green, N=37. The proportion of apolar (A), polar (B), or charged residues (C) was calculated for each protein. The average degree of solvent exposition was calculated using GETAREA. Exposition of apolar (D), polar (E), or charged (F) residues. Mann-Whitney test, **0.0013, ***<0.0001.

**Figure 4.**
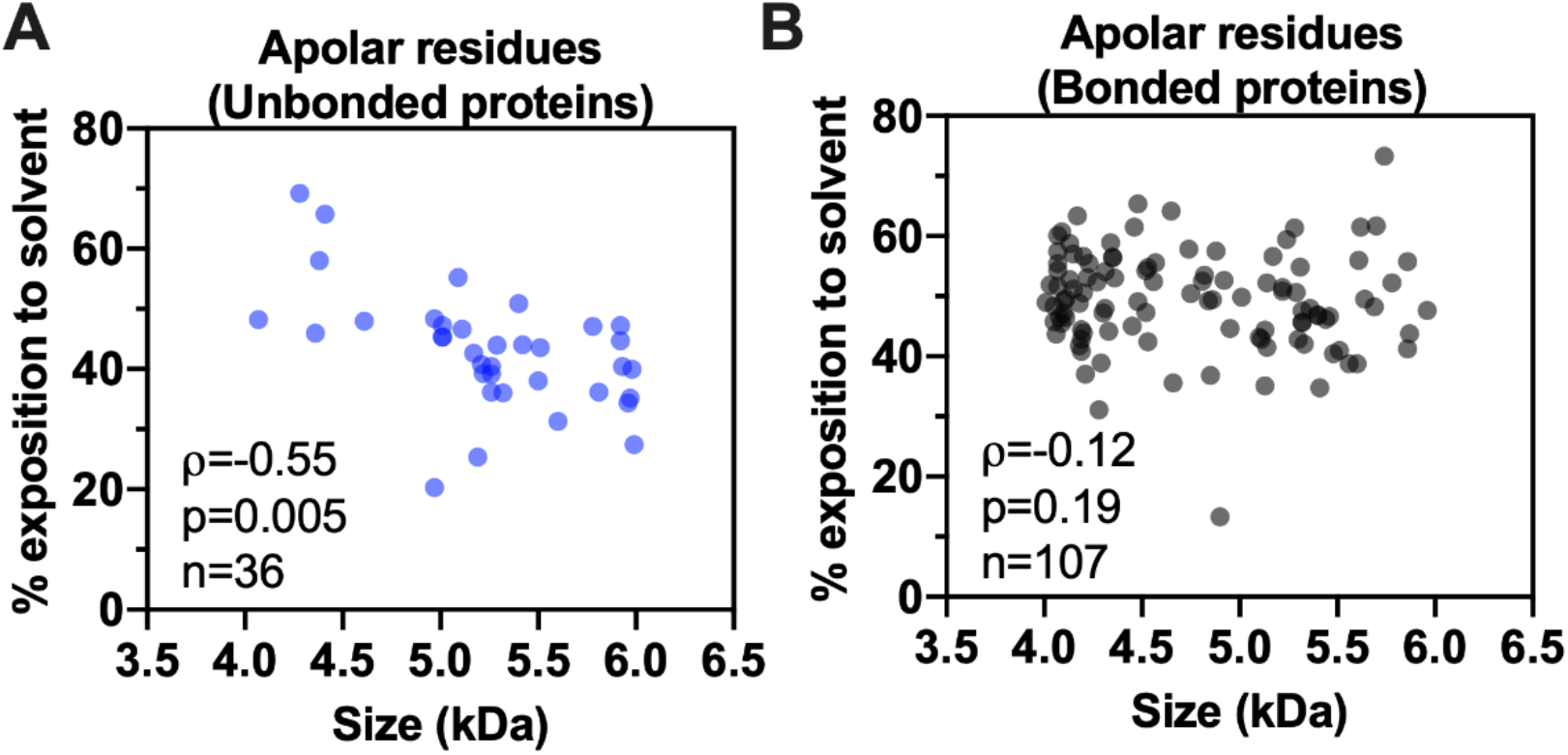
The unbonded group presents a good correlation between protein size and exposition to solvent for apolar residues. The Spearman correlation between apolar residues’ exposition with protein size for the unbonded group (A) or bonded group (B).

**Figure 5.**
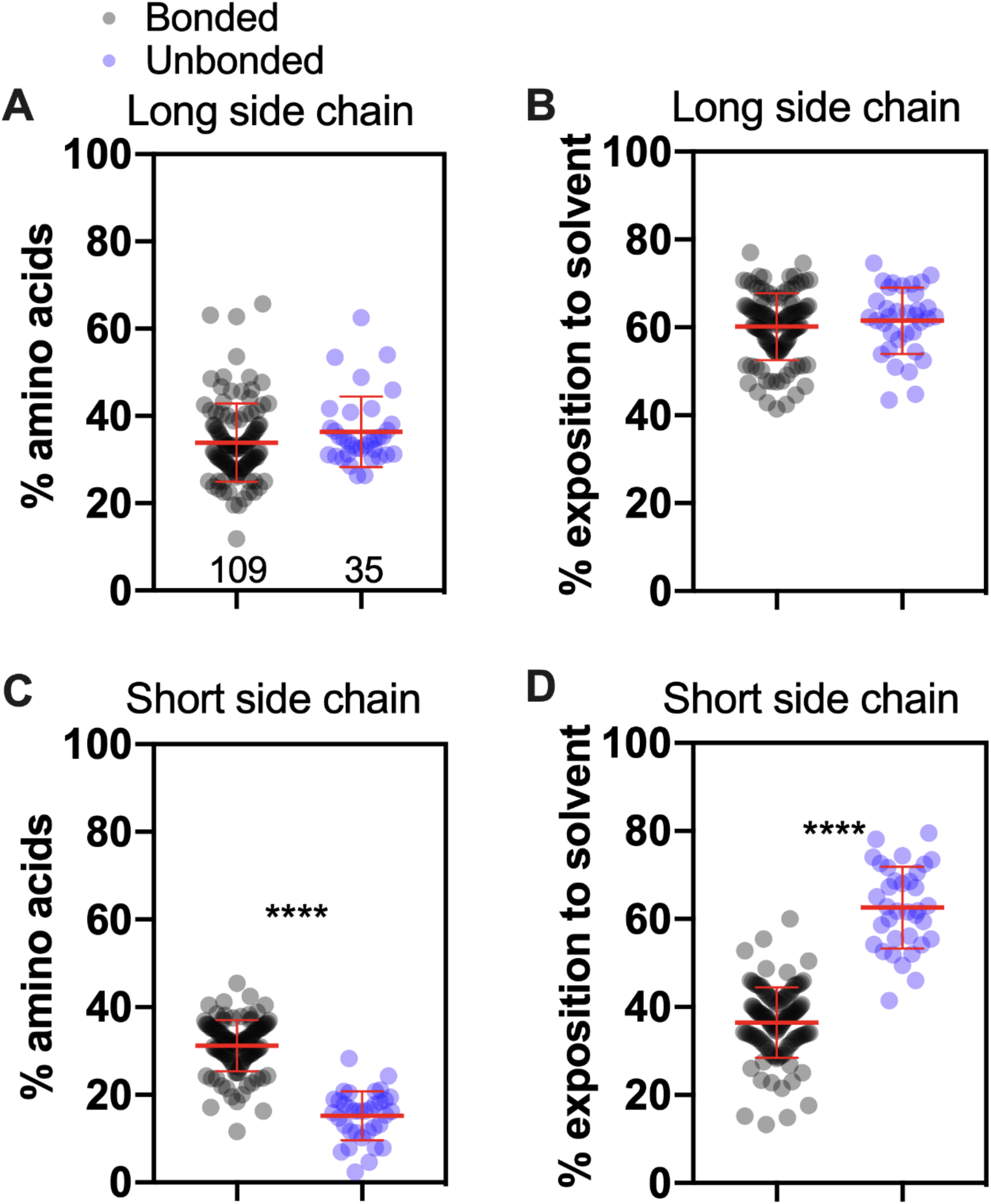
Comparison between proportion and exposition of long or short polar/charged residues for the bonded and unbonded group. In all graphs, the bonded group proteins are represented in black, and proteins of the unbonded group are in blue. Comparison of the proportion (A) and solvent exposition (B) of long-chain polar/charged (A) residues among the bonded vs. unbonded groups of proteins. The same analysis was performed for short-chain polar/charged residues (C and D). Mann-Whitney test, ****<0.0001.

We conclude that if the defensins do present an unusual fold (8, 9, 19), we can see that their structure is not unique compared to other similar small disulfide proteins.

### Small disulfide-rich proteins tend to expose more of their apolar residues to an aqueous solvent than non-disulfide proteins

Since no differences were observed within the bonded group (Figure 2), we then proceed to analyze how the bonded group as a whole differs from small proteins without disulfide bonds (unbonded group) following the same pipeline described in Figure 2. To define the unbonded group, we downloaded all PDB ranging from 4 to 6 kDa from RCSB PDB without disulfide bond (382 protein structures). We followed the same manual review described by the bonded group before, but this time we end up with just 36 proteins (9,4 % of the initial sample). It is important to note that even starting with about the same proteins, after manual revision, the group of bonded proteins end up with 3-fold more proteins. One explanation for this discrepancy may be because small proteins without disulfide bonds are more dependent on cofactors to achieve their fold. Figure 3 shows a comparison between bonded group vs. unbonded group for amino acid classification (apolar, polar, and charged), in terms of proportion within the primary sequence (Figure 3A-C), and their exposition to solvent (GETAREA analyzes, Figures 3 D-F). It is remarkable that essentially all parameters analyzed presented significant differences between these groups. As observed in Figures 3A and 3D, the bonded group, besides the lower proportion of apolar residues, can afford a higher degree of exposition to the aqueous solvent. This result leads us to conclude that this group of proteins does not present a canonical hydrophobic core in their structure. Even though it contradicts the expected folding model, it is according to what was observed for the structure of cysteine-rich peptides (7–9, 19).

Due to their tertiary structure stabilized by the disulfide bonds, proteins in the bonded group can afford this seemingly costly conformation. These proteins collapse in an enthalpic and entropic favorable ensemble, which allows the formation of disulfide bonds (6, 20). However, their hydrophobic residues’ exposition to the solvent makes these proteins more prone to form aggregates(6).

A similar conclusion was obtained analyzing the area of solvent accessibility through the Chimera 1.14 software (Figure S2). The result of Figures 3F (GETAREA analyzes) did not show a significant difference between groups regarding the exposition of charged residues. For the other side, when the structures were analyzed by Chimera 1.14, we observed a slightly higher exposition of charged residues for the unbonded group compared to the bonded group (Figure S2F).

We also analyzed whether exposing or hiding amino acid residues was correlated to the protein size. Even though there is no consensus in the folding models for proteins with disulfide bonds in their structure, it is suggested that it is due to their disulfide bonds that these proteins have a stable structure(21). Therefore, it is unlikely that the tertiary structure tendency observed for this group of small proteins would be affected by the length of their amino acid chains because of their stability. However, as the protein size increases, the folding pathway for the proteins of the unbonded group would be less affected by steric hindrance. Therefore a canonical hydrophobic collapse would be more likely to occur. With that being said, we calculated the correlation of the amount of exposition of apolar, polar, and charged residues vs. the protein size for both groups, unbonded and bonded. Only the unbonded group presents a negative correlation (ρ=−0.55) between protein size and exposition to solvent for apolar residues, as shown in Figure 4.

The result of Figure 4 reveals that even in a narrow size range (4 to 6 kDa), we can see the tendency of less exposition of the apolar residues as the protein size increases for proteins in the unbonded group. For example, the smallest protein in the group, PDB ID 1WY3, has an average exposition of 48% for its apolar residues, while for the largest protein, PDB ID 2N8O, this exposition drops to 27%. This does not occur for the bonded group, probably due to their more rigid structure, so it is more difficult for them to find other folded conformations. It is important to note that the bonded group proteins seem to not present their lowest free energy conformation as their native state, which is a possibility stated by Levinthal(22).

Pinheiro-Aguiar and colleagues noticed the presence of hydrophobic surface-clusters at defensin 2 from *Pisum sativum*(9). The authors suggested that long polar/charged side chains at the surface-clusters protect the hydrophobic amino acids from complete exposure to the solvent(9). The following experiment was designated to address if the bonded group has some preferences for long polar/charged amino acids compared to the unbonded group. The amino acids were classified as short polar/charged (CYS, SER, ASN, ASP) or long polar/charged (THR, TYR, GLN, LYS, ARG, HIS, GLU, see also Table 3). The remaining amino acids (GLY, ALA, VAL, PRO, LEU, ILE, MET, TRP, PHE) were omitted in this analyzes. First, the proportion in the primary sequence of long polar/charged residues was compared between bonded vs. unbonded groups. If the hypothesis of Pinheiro-Aguilar is correct, we expect to see a higher proportion of long polar/charged in the bonded group, but it was not the case (Figure 5A). No difference also was found comparing the degree of solvent exposition of long polar/charged residues between the groups (Figure 5B). For the other side, we observed differences for both the proportion and degree of solvent exposition of short polar/charged residues between the groups (Figures 5C-D). The bonded group presented a higher proportion and a lower exposition to the solvent of short polar/charged compared to the unbonded group (Figures 5C-D). We conclude that no bias toward long polar/charged residues exists for the bonded group. It is unlikely that the model proposed by Pinheiro-Aguiar and colleagues explains the unusual solvent exposition of hydrophobic residues that occurs in the fold of defensins and other small disulfide-rich proteins.

### Prediction algorithms have difficulty solving tertiary structures of small cysteine-rich proteins

To understand if the structure of the bonded group peptides presents a non-obvious folding, we challenged the predictor server Robetta selecting the *ab initio* option. We systematically submitted all proteins to the server with this condition because we wanted to compare how accurate the prediction is for both groups under the same conditions. Then, the structure obtained by Robetta was compared to the PDB file obtained from RCSB PDB, and the Chimera 1.14 software calculated the Root-mean-square deviation (RMSD) of atomic positions. Therefore, any remarked difference between the RMSDgroups’ RMSD values is due solely to the predicted folding based on the proteins’ primary sequences.

Figure 6A shows an example of accurate prediction: the original PDB (5DMA, gold) superposed well with the predicted structure by Robbeta (cyan) with an RMSD of one angstrom. The other scenario was observed for the PDB 2N2Q, where the RMSD obtained was eleven angstroms (Figure 6B). Figure 6C illustrates the comparison of RMSD values between the groups. We observed that the software Robbeta predicted with higher accuracy the proteins belonging to the unbonded group (Figure 6C). Finally, we correlated the proportion and degree of solvent exposition of apolar, polar, and charged residues with the RMSD. Again, we divided the protein into two groups, unbonded and bonded. Figure S3 shows the Spearman correlation for the parameter described before. We observed correlations with a p-value below 0.05 for two analyses, both for the unbonded group (Figure S3B). The amount of polar residues correlates positively with the RMSD ((ρ=0,43, Figure S3C). The degree of solvent exposition of apolar residues also presented a positive correlation (ρ=0.51) with RMSD. The Rosetta all-atom forcefield is based on hydrogen bonding, short-range Van der Waals interactions, and desolvation(18, 23). Although very realistic, this forcefield did not predict well the structure of our groups of peptides, especially those with disulfide bonds. This could indicate that the native conformation of the bonded group proteins is not their lowest free-energy state, and considering the high degree of their apolar residues’ exposition, this lack of hydrophobic core is probably not an evident folding. The structural prediction of small proteins is a challenging task(24), which is improving due to deep learning techniques(25), but still needs development, especially for defensin and small disulfide-rich proteins.

**Figure 6.**
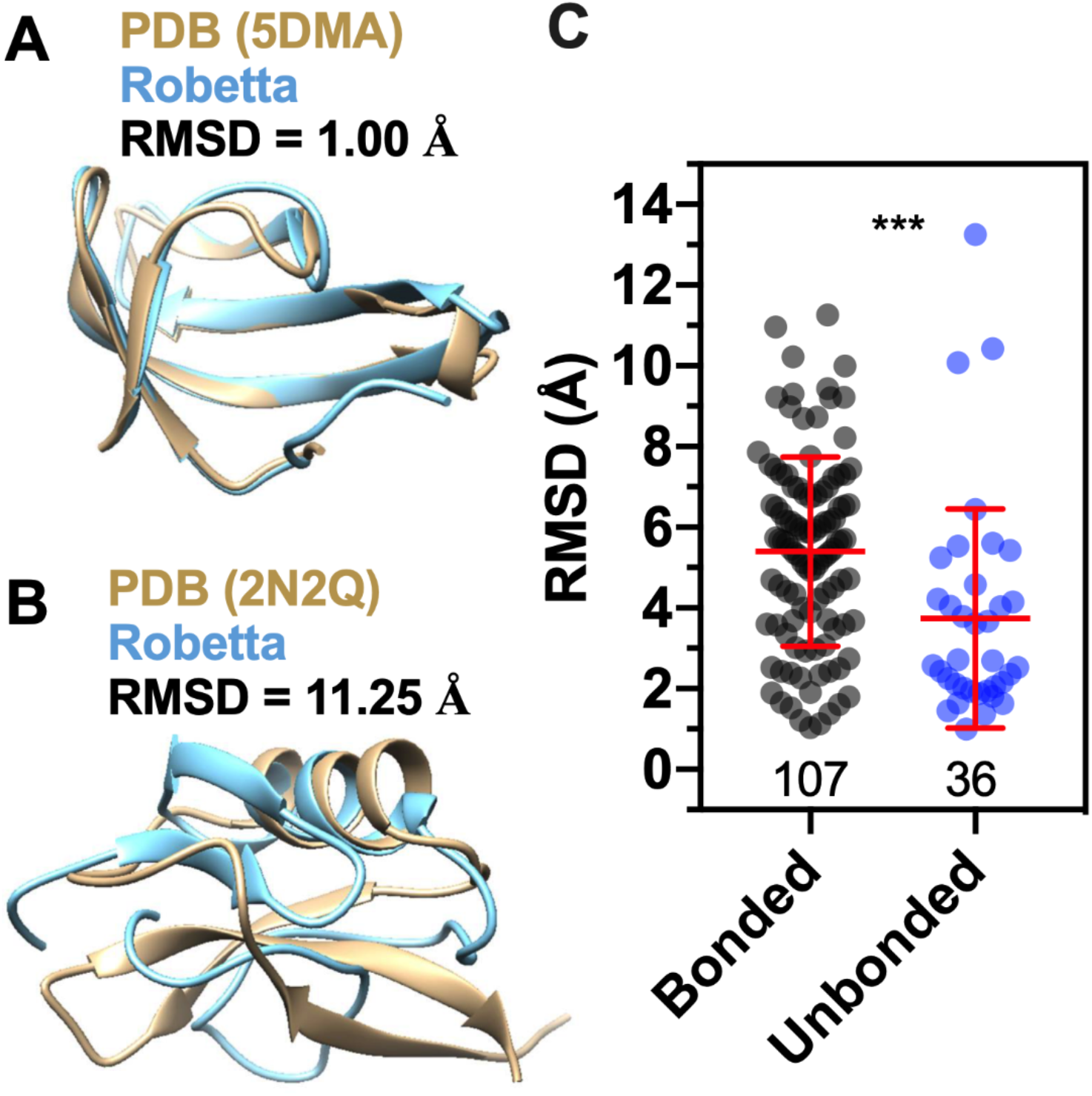
The prediction accuracy of the *ab initio* algorithm Robetta was higher for the unbonded when compared to bonded proteins. Visual representation between original PDB file for 5DMA (A) or 2N2Q (B) with the structure predicted by Robetta. (C) Comparison between RMSD means for the bonded and unbonded groups. Mann-Whitney test, ***<0.0001.

## Supporting information

Support Information (Table S1)

## Data availability

The algorithms used in this work are available in the GitHub repository (HTTPS://github.com/mhoyerm).

## Supplementary data

Supplemental data for this article can be accessed on the publisher’s website.

## Acknowledgments

We thank Ramon Pinheiro-Aguiar for helpful discussions.

## Abbreviations

PDB: Protein Data Bank
RMSD: root-mean-square deviation

---

**Figure S1.**
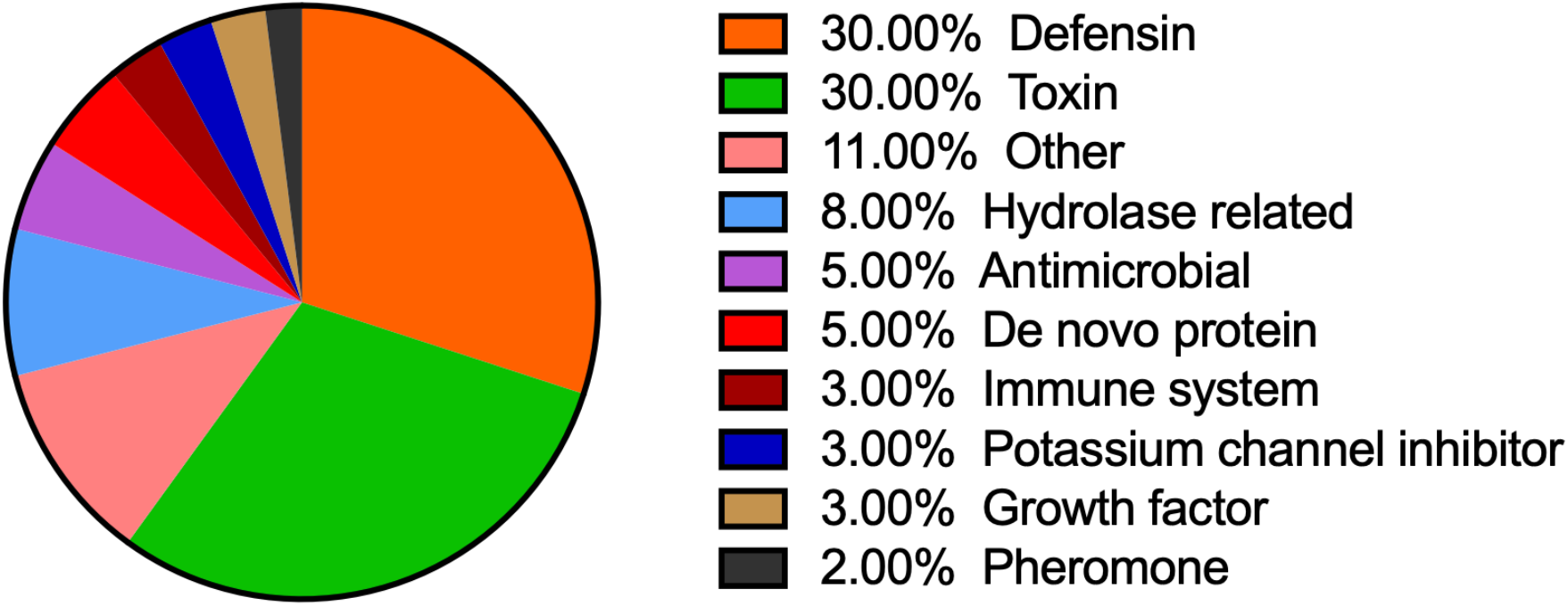
Classification and proportion of proteins in the bonded group. Defensin and defensin-like proteins represent 30% of the proteins from 108 proteins belonging to the bonded group.

**Figure S2.**
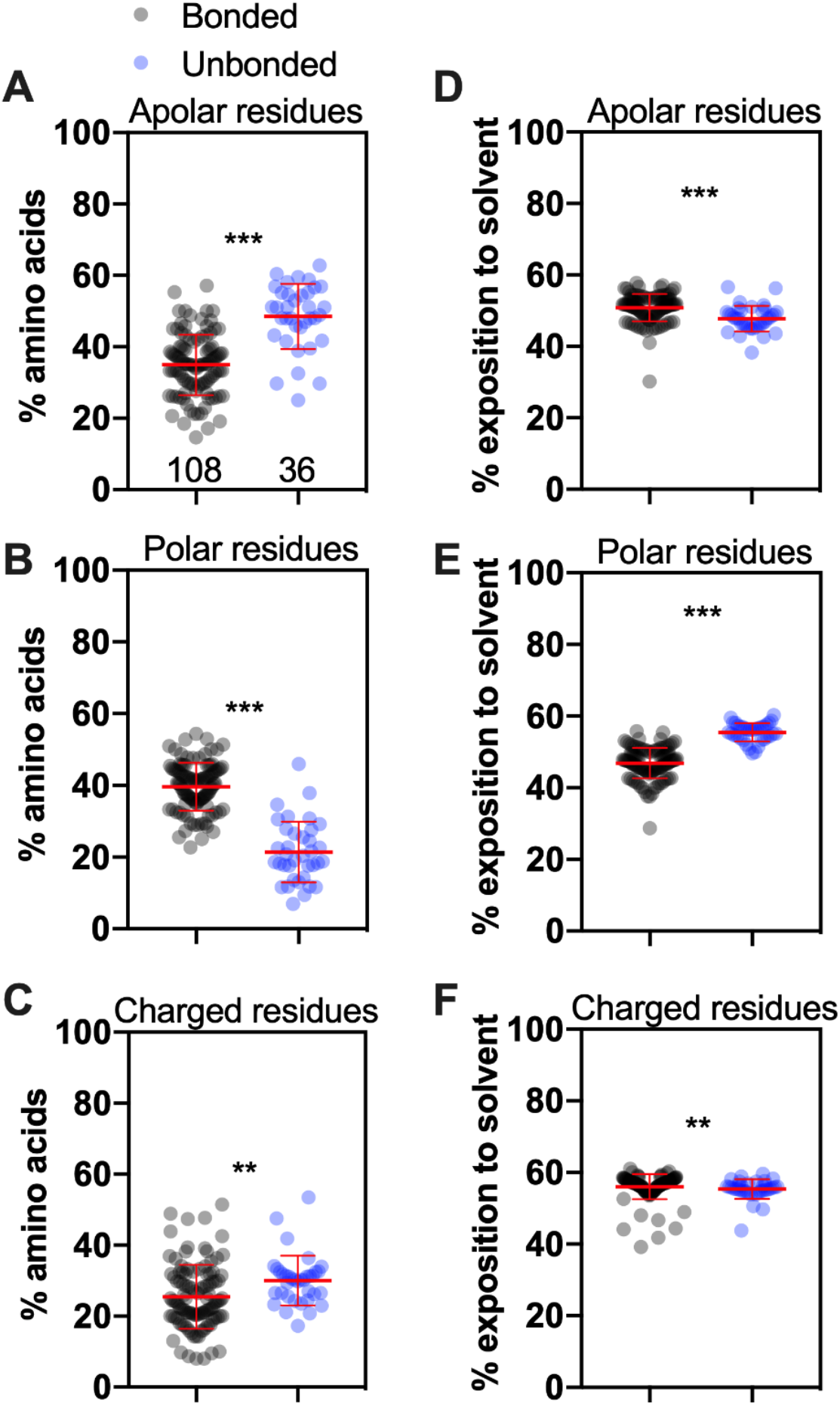
Small proteins with at least one disulfide bond (bonded) present distinct features regarding the proportion and exposition of residues compared to proteins that do not make disulfide bonds (unbonded). In all graphs, the whole population of bonded proteins is represented in black, N=108, while the group of unbonded proteins is represented in green, N=37. The proportion of apolar (A), polar (B), or charged residues (C) was calculated for each protein. The average degree of solvent exposition was calculated using Chimera 1.14 software. Exposition of apolar (D), polar (E), or charged (F) residues. Mann-Whitney test, **0.0072, ***<0.0001.

**Figure S3.**
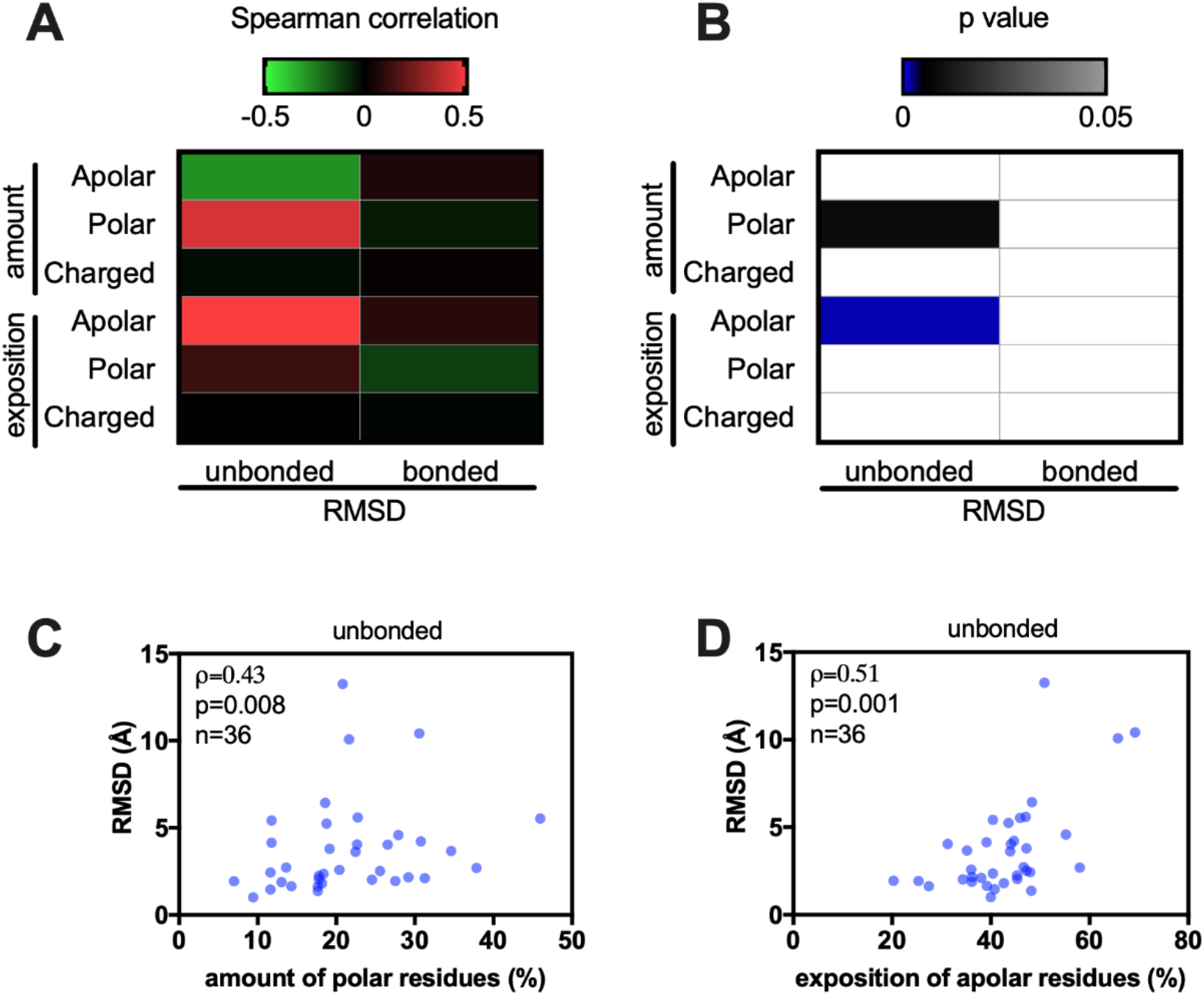
Spearman correlation of RMSD vs the amount or exposition of different classes of amino acids in bonded and unbounded groups. Heat map of the Spearman correlation (A) or p value (B) of RMSD calculated in Figure 6 (se also Table S1) and the amount or exposition of apolar, polar and charged residues. The proteins were divided in two groups, unbonded and bonded. Statistical significance (p<0.05) were observed for two comparisons, namely, RMSD vs amount of polar residues (C) and RMSD vs exposition of apolar residues (D), both for the unbonded group.

## References

1. Shafee,T.M.A., Lay,F.T., Phan,T.K., Anderson,M.A. and Hulett,M.D. (2017) Convergent evolution of defensin sequence, structure and function. Cell Mol Life Sci, 74, 663–682.

2. Shafee,T.M.A., Lay,F.T., Hulett,M.D. and Anderson,M.A. (2016) The Defensins Consist of Two Independent, Convergent Protein Superfamilies. Molecular Biology and Evolution, 33, 2345–2356.

3. Mookherjee,N., Anderson,M.A., Haagsman,H.P. and Davidson,D.J. (2020) Antimicrobial host defence peptides: functions and clinical potential. Nat Rev Drug Discov, 19, 311–332.

4. Almeida,M.S., Cabral,K.M.S., Kurtenbach,E., Almeida,F.C.L. and Valente,A.P. (2002) Solution structure of Pisum sativum defensin 1 by high resolution NMR: plant defensins, identical backbone with different mechanisms of action. J. Mol. Biol., 315, 749–757.

5. Undheim,E.A.B., Mobli,M. and King,G.F. (2016) Toxin structures as evolutionary tools: Using conserved 3D folds to study the evolution of rapidly evolving peptides. BioEssays, 38, 539–548.

6. Fraga,H., Graña-Montes,R., Illa,R., Covaleda,G. and Ventura,S. (2014) Association between foldability and aggregation propensity in small disulfide-rich proteins. Antioxid Redox Signal, 21, 368–383.

7. Rose,G.D., Geselowitz,A.R., Lesser,G.J., Lee,R.H. and Zehfus,M.H. (1985) Hydrophobicity of amino acid residues in globular proteins. Science, 229, 834–838.

8. Machado, LESF, De Paula,V.S., Pustovalova,Y., Bezsonova,I., Valente,A.P., Korzhnev,D.M. and Almeida,F.C.L. (2018) Conformational Dynamics of a Cysteine-Stabilized Plant Defensin Reveals an Evolutionary Mechanism to Expose Hydrophobic Residues. Biochemistry, 57, 5797–5806.

9. Pinheiro-Aguiar,R., do Amaral,V.S.G., Pereira,I.B., Kurtenbach,E. and Almeida,F.C.L. (2020) Nuclear magnetic resonance solution structure of Pisum sativum defensin 2 provides evidence for the presence of hydrophobic surface-clusters. Proteins, 88, 242–246.

10. Cheng,J., Choe,M.-H., Elofsson,A., Han,K.-S., Hou,J., Maghrabi,A.H.A., McGuffin,L.J., Menéndez-Hurtado,D., Olechnovič,K., Schwede,T., et al. (2019) Estimation of model accuracy in CASP13. Proteins, 87, 1361–1377.

11. Service, RF (2020) ‘The game has changed.’ AI triumphs at protein folding. Science, 370, 1144–1145.

12. Dill,K.A. and MacCallum,J.L. (2012) The protein-folding problem, 50 years on. Science, 338, 1042–1046.

13. AlQuraishi,M. (2019) End-to-End Differentiable Learning of Protein Structure. Cell Systems, 8, 292–301.e3.

14. Fraczkiewicz,R. and Braun,W. (1998) Exact and efficient analytical calculation of the accessible surface areas and their gradients for macromolecules. Journal of Computational Chemistry, 19, 319–333.

15. Pettersen,E.F., Goddard,T.D., Huang,C.C., Couch,G.S., Greenblatt,D.M., Meng,E.C. and Ferrin,T.E. (2004) UCSF Chimera—A visualization system for exploratory research and analysis. Journal of Computational Chemistry, 25, 1605–1612.

16. Sanner,M.F., Olson,A.J. and Spehner,J.C. (1996) Reduced surface: An efficient way to compute molecular surfaces. Biopolymers, 38, 305–320.

17. Song,Y., DiMaio,F., Wang,R.Y.-R., Kim,D., Miles,C., Brunette,T., Thompson,J. and Baker,D. (2013) High-resolution comparative modeling with RosettaCM. Structure, 21, 1735–1742.

18. Raman,S., Vernon,R., Thompson,J., Tyka,M., Sadreyev,R., Pei,J., Kim,D., Kellogg,E., DiMaio,F., Lange,O., et al. (2009) Structure prediction for CASP8 with all-atom refinement using Rosetta. Proteins, 77, 89–99.

19. Torres,A.M., de Plater,G.M., Doverskog,M., Birinyi-Strachan,L.C., Nicholson,G.M., Gallagher,C.H. and Kuchel,P.W. (2000) Defensin-like peptide-2 from platypus venom: member of a class of peptides with a distinct structural fold. Biochem J, 348 Pt 3, 649–656.

20. Narayan,M. (2020) Revisiting the Formation of a Native Disulfide Bond: Consequences for Protein Regeneration and Beyond. Molecules 2020, Vol. 25, Page 5337, 25, 5337.

21. Robinson,P.J. and Bulleid, NJ (2020) Mechanisms of Disulfide Bond Formation in Nascent Polypeptides Entering the Secretory Pathway. Cells 2020, Vol. 9, Page 1994, 9, 1994.

22. Levinthal,C. (1968) Are there pathways for protein folding? J. Chim. Phys., 65, 44–45.

23. Protein Structure Prediction Using Rosetta (2004) Protein Structure Prediction Using Rosetta. Methods in Enzymology, 383, 66–93.

24. Bradley,P., Misura, KMS and Baker,D. (2005) Toward high-resolution de novo structure prediction for small proteins. Science, 309, 1868–1871.

25. Senior,A.W., Evans,R., Jumper,J., Kirkpatrick,J., Sifre,L., Green,T., Qin,C., Žídek,A., Nelson,A.W.R., Bridgland,A., et al. (2020) Improved protein structure prediction using potentials from deep learning. Nature, 577, 706–710.

